# Volume-wise analysis of fMRI time series allows accurate prediction of language lateralization

**DOI:** 10.1101/725671

**Authors:** Martin Wegrzyn, Markus Mertens, Christian G. Bien, Friedrich G. Woermann, Kirsten Labudda

**Affiliations:** Department of Psychology, Bielefeld University, Bielefeld, Germany; Bethel Epilepsy Center, Krankenhaus Mara, Bielefeld, Germany

**Keywords:** language lateralization, Wada test, pattern analysis, predictive modeling, functional MRI

## Abstract

Using fMRI as a clinical tool, for example for lateralizing language, requires that it provides accurate results on the individual level. However, using a single voxel-wise activity map per patient limits how well the uncertainty associated with a decision can be estimated. Here, we explored how using a “volume-wise” analysis, where the lateralization of each time point of a patient’s fMRI session is evaluated independently, could support clinical decision making. Ninety-six patients with epilepsy who performed a language fMRI were analyzed retrospectively. Results from Wada testing were used as an indication of true language lateralization. Each patient’s 200 fMRI volumes were correlated with an independent template of prototypical lateralization. Depending on the strength of correlation with the template, each volume was classified as indicating either left-lateralized, bilateral or right-lateralized language. A decision about the patient’s language lateralization was then made based on how most volumes were classified. The results show that, using a simple majority vote, accuracies of 84% were reached in a sample of 63 patients with high-quality data. When 33 patients with datasets previously deemed inconclusive were added, the same accuracy was reached when more than 43% of a patient’s volumes were in agreement with each other. Increasing this cutoff to 51% volumes with agreeing classifications allowed for excluding all inconclusive cases and reaching accuracies over 90% for the remaining cases. Further increasing the cutoff to 65% agreeing volumes resulted in correct predictions for all remaining patients. The study confirms the usefulness of fMRI for language lateralization in patients with epilepsy, by demonstrating high accuracies. Furthermore, it illustrates how the diagnostic yield of individual volumes of fMRI data can be increased using simple similarity measures. The accuracy of our approach increased with the number of agreeing volumes, and thus allowed estimating the uncertainty associated with each individual diagnosis.

## Introduction

For functional MRI to be clinically useful, it has to work on the individual level (Gabrieli et al., 2015; Woo et al., 2017). However, translating the knowledge generated from fMRI studies into clinical applications has proven challenging (Bandettini, 2018). A single fMRI session, lasting only a few minutes, provides rich information in space (usually tens of thousands of voxels) and time (usually hundreds of volumes). It would seem that this wealth of data should allow performing in-depth analyses on the level of N=1 (Smith and Little, 2018). Meanwhile, neuroscientists are questioning whether sample sizes as large as N=100 are sufficient to generate replicable fMRI results (Turner et al., 2018). One reason for this schism might be the framework of voxel-wise analysis, where each voxel is treated as independent of the rest of the brain. Because this approach was not developed with clinical applications in mind, it has been argued that using fMRI as a biomarker requires a shift in perspective (Woo et al., 2017; Kragel et al., 2018).

To capitalize on the rich spatial data that fMRI provides even in the individual case, pattern analysis methods try to aggregate information across the whole brain. Distributed patterns of brain activity can function as fingerprints, unique to different tasks or persons (Haxby, 2001; Finn et al., 2015). This approach can provide the power needed to find effects on the individual level, by turning miniscule voxel-wise differences into large differences of pattern dissimilarity (Woo et al., 2017). Finally, the aggregation of all information across the brain into one prediction seems useful when clinical decisions have to be made (Woo et al., 2017).

One application where fMRI is successfully used as a diagnostic tool is the lateralization of language functions (Bradshaw et al., 2017b; Szaflarski et al., 2017). Patients with epilepsy who do not respond to a pharmacotherapy sometimes have to undergo surgery in order to become seizure-free. To plan the resection in a way that cortex critical for a certain cognitive function is spared, a number of presurgical tests needs to be performed. As language functions are critical for everyday life and strongly lateralized to one side of the brain (Frost, 1999), resection of eloquent cortex in the dominant hemisphere needs to be avoided. Identifying language dominance is especially important in epilepsy, since the incidence of atypical (bilateral or right-lateralized) language in patients is as high as 20% (Springer et al., 1999).

The current gold standard for language lateralization is the invasive Wada test, which temporarily disrupts the function of one hemisphere to evaluate the language abilities of the other hemisphere in isolation (Kurthen et al., 1994). Although fMRI measures only activity and cannot simulate a functional loss, both methods have a high level of agreement (Binder et al., 1996; Adcock et al., 2003; Woermann et al., 2003; Dym et al., 2011; Janecek et al., 2013). Usually, the fMRI activity map for a language task, as compared to a control condition, is computed using a voxel-wise approach. Then, a laterality index is computed, for example by counting how many voxels fall above a certain threshold in the left and right frontal lobe (Wilke and Lidzba, 2007; Bradshaw et al., 2017a). While the advantages of different implementations of the laterality index are under debate (Wilke and Lidzba, 2007; Bradshaw et al., 2017a), they have proven to be robust, provided that the data are of sufficient quality (Wegrzyn et al., 2019).

With fMRI being able to predict the results of the Wada test with accuracies around 90% (Dym et al., 2011; Bauer et al., 2014) it has become an established tool for lateralizing language (Binder, 2011; Szaflarski et al., 2017; Benjamin et al., 2018). Its non-invasive nature is an important asset for a biomarker, as it can be easily repeated, extended and used on many patients, as well as healthy normative samples (Gabrieli et al., 2015; Dubois and Adolphs, 2016). However, there are a number of challenges yet to be solved. A number of studies have shown that fMRI is able to predict typical left-lateralization with high accuracy, but it is less reliable for atypical cases (Benke et al., 2006; Arora et al., 2009; Janecek et al., 2013; Benjamin et al., 2018). One meta-analysis showed accuracies around 95% for typical and only around 50% for atypical cases (Bauer et al., 2014). This led the authors to conclude that fMRI should only be trusted when it indicates left-lateralization. A second problem is that the common ways to compute a laterality index critically rely on data pre-selection (Wilke and Lidzba, 2007; Benjamin et al., 2017). Depending on the method used, a dataset comprised of only noise can easily produce a laterality index indicative of strong lateralization or of perfect bilaterality (Wegrzyn et al., 2019). Therefore, an outcome that reflects a measure of confidence in its reliability would be desirable. On the group level, the accuracy of a method is determined by the number of correctly diagnosed patients. Given that each fMRI dataset is comprised not only of many voxels in space, but also of many observations in time, similar accuracy estimates could be attempted on the level of N=1. This would mean that each volume of a patient’s fMRI dataset would have to be diagnosed separately and the overall agreement between volumes could serve as a measure of uncertainty.

Previous work indicates that this could be feasible, given that language processing can be distinguished from other tasks with near-perfect accuracies using short blocks of an individual person’s fMRI (Wegrzyn et al., 2018). In the present work, we further explore how shifting from a “voxel-wise” to a “volume-wise” analysis could complement established methods of language lateralization. The present study used language fMRI data from patients with epilepsy who also underwent Wada testing. Each patient’s fMRI session consisted of 200 volumes, which were classified separately as being indicative of left-lateralized, bilateral or right-lateralized language. This decision depended on how strongly the left-right asymmetry of each volume correlated with an independent template of typical lateralization. Based on how many of a patient’s volumes were classified the same way, a probabilistic decision about that patient’s language lateralization was made.

## Methods

### 2.1 Participants

The study included 96 patients with epilepsy who were undergoing a presurgical evaluation with fMRI and Wada testing. All patients performed an fMRI verbal-fluency task at the Epilepsy Center Bethel between September 2011 and March 2018. The start of the inclusion period was determined by the installation of a 3T scanner at the study site. All data were acquired as part of the center’s presurgical evaluation program, and were analyzed retrospectively. The study was approved by the ethics board of Bielefeld University (2017-184). Thirty-seven of the patients were female (39%) and the median age was 24 years (range 7-61).

### 2.2 fMRI data acquisition

Data were collected on a 3T Siemens Verio MR scanner. The fMRI data were collected using a 12-channel head coil using the following parameters: 21 axial slices per volume, 3×3 mm in-plane resolution for each slice and a thickness of 5 mm. A repetition time (TR) of 3 seconds was used with an effective acquisition time of 1.8 seconds per volume; there was a 1.2-second pause between TRs to allow for the audible presentation of verbal instructions. Two hundred volumes, not counting two dummy scans, were collected over a period of ten minutes.

### 2.3 fMRI task

Each patient performed a verbal fluency task, which consisted of covert word production of either semantic or phonemic categories, such as “animals” or words beginning with the letter “S”. Whether a patient received the semantic or the phonemic task was decided individually, taking into account the patient’s abilities and experience with the task. A block of verbal fluency lasted 30 seconds, and was alternated with a resting block of equal length. There were 10 task and 10 rest blocks, each of which was triggered by a verbal instruction given via the MRI intercom. A sample dataset, including prototypical instructions for the semantic task, can be found on OpenNeuro (10.18112/openneuro.ds002014.v1.0.1).

### 2.4 fMRI data preprocessing

Minimal preprocessing was performed using SPM12 (fil.ion.ucl.ac.uk/spm). After movement correction using the realign function, the images were normalized to the echoplanar imaging (EPI) template and up-sampled to 2 mm isotropic voxels (Calhoun et al., 2017). Further preprocessing and transformation of data was performed using nilearn 0.5.0 as implemented in Python 3.6. Data were smoothed with 6 mm FWHM and the time course of each voxel was z-scored to have zero mean and unit variance. Linear drifts and movement parameters derived from the realignment were regressed out as confounds.

### 2.5 Pre-selection of data

The 96 datasets used in the main analysis were not pre-selected based on any criteria of quality or task performance. This was important as the aim of the present study was to construct a measure of language lateralization which would reflect the uncertainty associated with each decision and not depend on any additional intervention. However, while all data were used for the final testing of our model’s performance (cf. section 2.8), a pre-selection of data was performed for the initial training of the classifier (cf. section 2.7). In this way, the classifier was allowed to learn how a prototypical conclusive fMRI looked like and later assign higher confidence to those cases. To keep training and test data separate, a leave-one-patient-out cross-validation scheme was employed (Esterman et al., 2010). Therefore, there were as many training folds as there were pre-selected patients, and each patient’s language lateralization was predicted using a model trained on independent patients. The data were pre-selected using the Python module described in Wegrzyn et al. (2019) and available on github.com/mwegrzyn/laterality-index-deconstruction. In brief, this procedure uses a whole-brain map of t-values and estimates the overall conclusiveness of the dataset by considering the numerator (L-R) and the denominator (L+R) of the common laterality index (LI) as two separate features. If these analyses deemed the dataset inconclusive, it was held back from training and only used later during testing of the classifier’s performance. Of the 96 fMRI datasets in the sample, 63 were preselected as conclusive. The activity maps generated for this step were also used for initial visual inspection of data in the canonical voxel-wise fashion (result section 3.1).

### 2.6 Feature extraction

To extract a measure of lateralization, each volume was flipped along the x-axis and the resulting mirror image subtracted from the original image. The resulting map indicated the left-right difference of each voxel with its contralateral homologue (Fig. 1, upper row). Each of these difference maps was then correlated with a template, which was also a left-right difference map representing typical left-lateralization (Fig. 1, left hand). This template, based on a sample of 382 independent patients, was taken from Wegrzyn et al. (2019), where its creation is described in detail. It is also available from NeuroVault (Gorgolewski et al., 2015): identifiers.org/neurovault.image:113673. A Pearson correlation was computed between each volume of a patient’s time course and the template (Fig. 1, middle row), resulting in 200 correlations per patient. A positive correlation with the template during task performance would indicate stronger left-lateralization and a negative correlation would indicate stronger right-lateralization. A weak correlation around zero should indicate bilaterality. During rest, the relationship between correlations and lateralization should be inverted. As the data were based on z-scored time courses of zero mean, positive values at one time point will be balanced by negative values at another time point. Therefore, we were able to treat task and rest periods as equally informative, and only inverted the sign of the correlations during periods of rest. Of note, this also means that after z-transformation the volumes are not independent of each other: Each volume can be thought of as carrying some information about the language-rest contrast, which can only be systematically positive during task periods if it is also systematically negative during rest periods. To account for the delay in the hemodynamic response function (HRF), the onsets of each block were shifted by two volumes (6 seconds) with respect of the actual timing of events. Such a simple boxcar function does not take into account the shape of the HRF, but has been shown to be effective (Wegrzyn et al., 2018). Not weighting each volume by its expected response was also important because the subsequent steps of training and testing the classifier were designed to treat all 200 volumes as independent observations of equal importance. The volume-wise correlations generated in this step were also used for initial visual inspection of data (result section 3.2).

**Figure 1.**
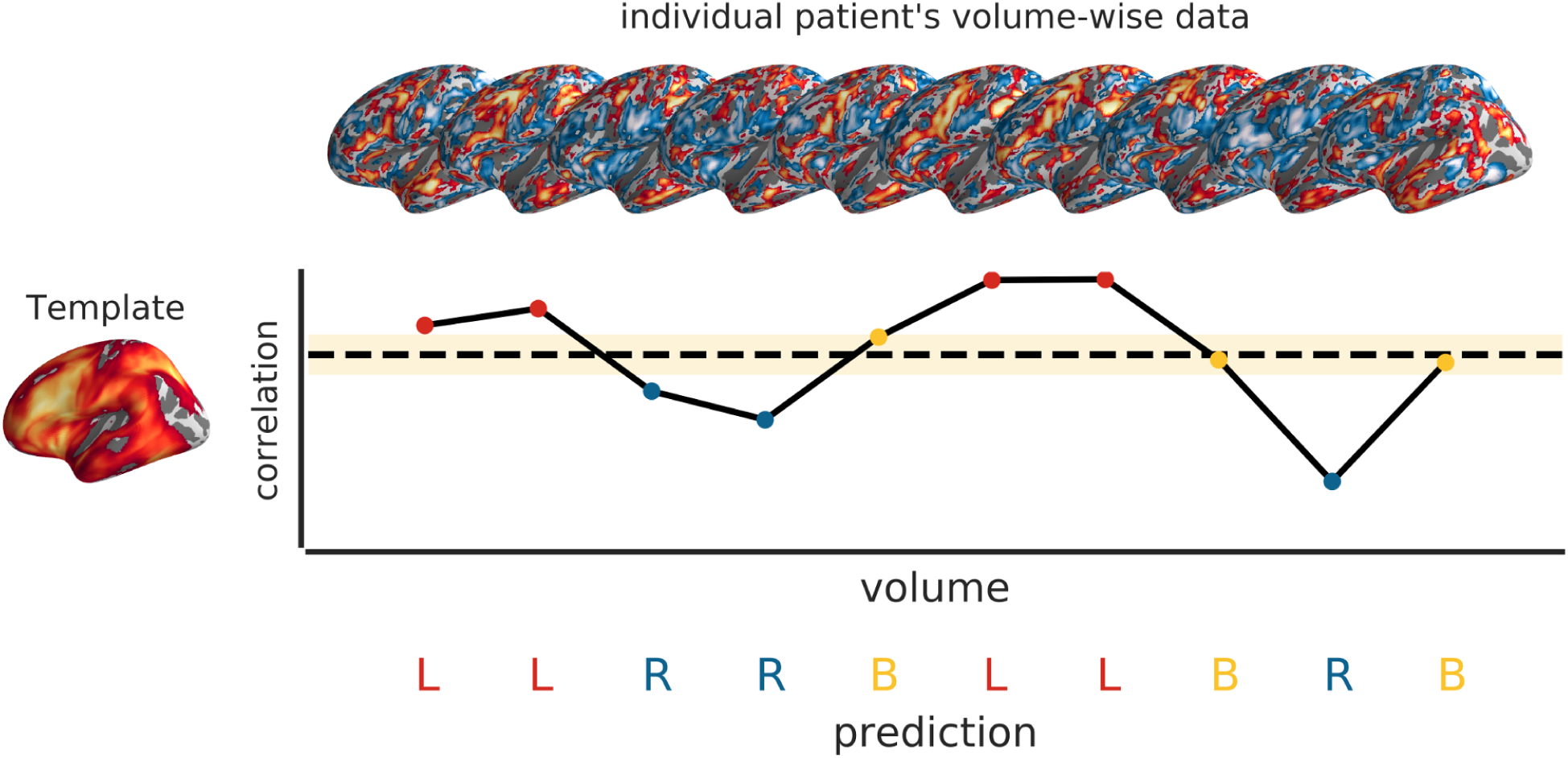
Illustration of making volume-wise predictions. In the upper row of the figure, ten brain volumes from one task block of one patient are shown. Each volume shows the left-right difference of its z-scored voxels. In the figure, the volumes are thresholded for illustrative purposes, but for analysis, unthresholded data were used. On the left-hand of the figure, a template of prototypical left-right asymmetry for the task is shown, based on data from Wegrzyn et al. (2019). Each individual volume can be correlated with this template. The resulting correlations are shown in the line plot in the middle of the figure. During a task block, more positive correlations with the template indicate stronger left-lateralization, while more negative correlations indicate stronger right-lateralization. An in-between range of values (beige area) around the dotted line at zero would be indicative of bilaterality. During rest blocks, the correlations would be inverted, so that right-lateralization during rest would be more indicative of left-lateralized language. What ranges of correlations should be classified as which type of lateralization would be determined by boundaries derived from a support-vector-classifier. The resulting volume-wise predictions are shown in the lower part of the figure (L=left; B=bilateral; R=right).

### 2.7 Training of the classifier

The goal of training was to determine which ranges of correlations were most indicative of left-lateralized, bilateral or right-lateralized language. The 200 volume-template correlations (cf. section 2.6) of all preselected patients (cf. section 2.5) were used to train a linear support vector classifier (SVC) as implemented in scikit-learn 0.19.1 (Pedregosa et al., 2011). Balanced class weights were used, so that correctly classifying the smaller samples of atypical cases (bilateral and right-lateralized) received equal weight to correctly classifying the typical left-lateralized cases. Each volume had one feature (correlation value) and one label (type of language lateralization).

The language lateralization of the volume (left-lateralized, bilateral or right-lateralized) was determined by the clinical evaluation of the respective patient’s Wada test result (Woermann et al., 2003; Wegrzyn et al., 2019). The categorical evaluation of the Wada test was taken from the patient’s medical records. Any Wada test with a categorical clinical assessment was included in the sample. Of the 96 patients, 59 were left-lateralized (61%), 14 were bilateral (15%) and 23 were right-lateralized (24%), according to the Wada test. The volumes were all treated as independent observations, so the total number of observations was n patients x 200 volumes. After training with a leave-one-patient-out cross-validation scheme, the boundaries (Fig. 1, middle row) of the classifiers were applied to each of the 200 volume-template correlations of the held-out patient. Depending on the range of values into which each correlation fell, it was classified as either indicating left-lateralized, bilateral or right-lateralized language. The middle row in Fig. 1 shows how each correlation within the beige-colored area would be classified as bilateral, while stronger positive correlations would be classified as left-lateralized and weaker negative correlations as right-lateralized. Finally, the patient’s overall language lateralization was decided by a winner-take-all classification, counting which type of lateralization was predicted by the relative majority of volumes.

### 2.8 Evaluation of predictions

The accuracy of the predictions was evaluated on the group level, by counting how many patients were correctly evaluated using the majority vote of their 200 volumes. Firstly, the accuracies were tested when only considering the pre-selected conclusive cases, as this scenario might allow closer comparability with the existing literature. Secondly, the accuracies were tested when considering all cases, even the ones who were previously excluded as inconclusive. This might allow better estimation of how the method would perform in a clinical context, when used on its own. In addition, the accuracies were tested when applying different cutoffs regarding the minimal amount of agreeing volumes. That is, only patients for whom the majority vote of their volumes exceeded a certain absolute number were considered. If the employed method reflected a measure of confidence on the individual level, accuracies should increase as the number of volumes with agreeing predictions increases.

The code used for training and testing the classifiers and to reproduce all figures is available on GitHub (github.com/mwegrzyn/volume-wise-language). The main modules used were numpy, scipy, scikit-learn, nilearn, matplotlib, seaborn, pycortex and jupyter.

## Results

### 3.1 Voxel-wise group analyses

The activity for the verbal fluency task compared to the rest condition for the patients in the training sample (N=63) is shown in Fig. 2. To allow for better comparability between the differently sized groups, the alpha-level was adjusted to keep power constant at 1-beta=0.9 for an expected effect size of d=0.8. Therefore, the smaller groups were thresh-olded at more liberal levels. All groups showed activity in a network of language- and task-related brain areas, including inferior and medial frontal areas, insula, supplementary motor area and intraparietal sulcus. Compared to the language task, the rest condition activated a characteristic default mode network, including precuneus, medial frontal and lateral parietal areas. This was true for all three groups. The same analyses based on the left-right difference images showed that the most lateralizing information was present in inferior and medial frontal areas. For the bilateral group, no clear lateralization pattern emerged.

**Figure 2.**
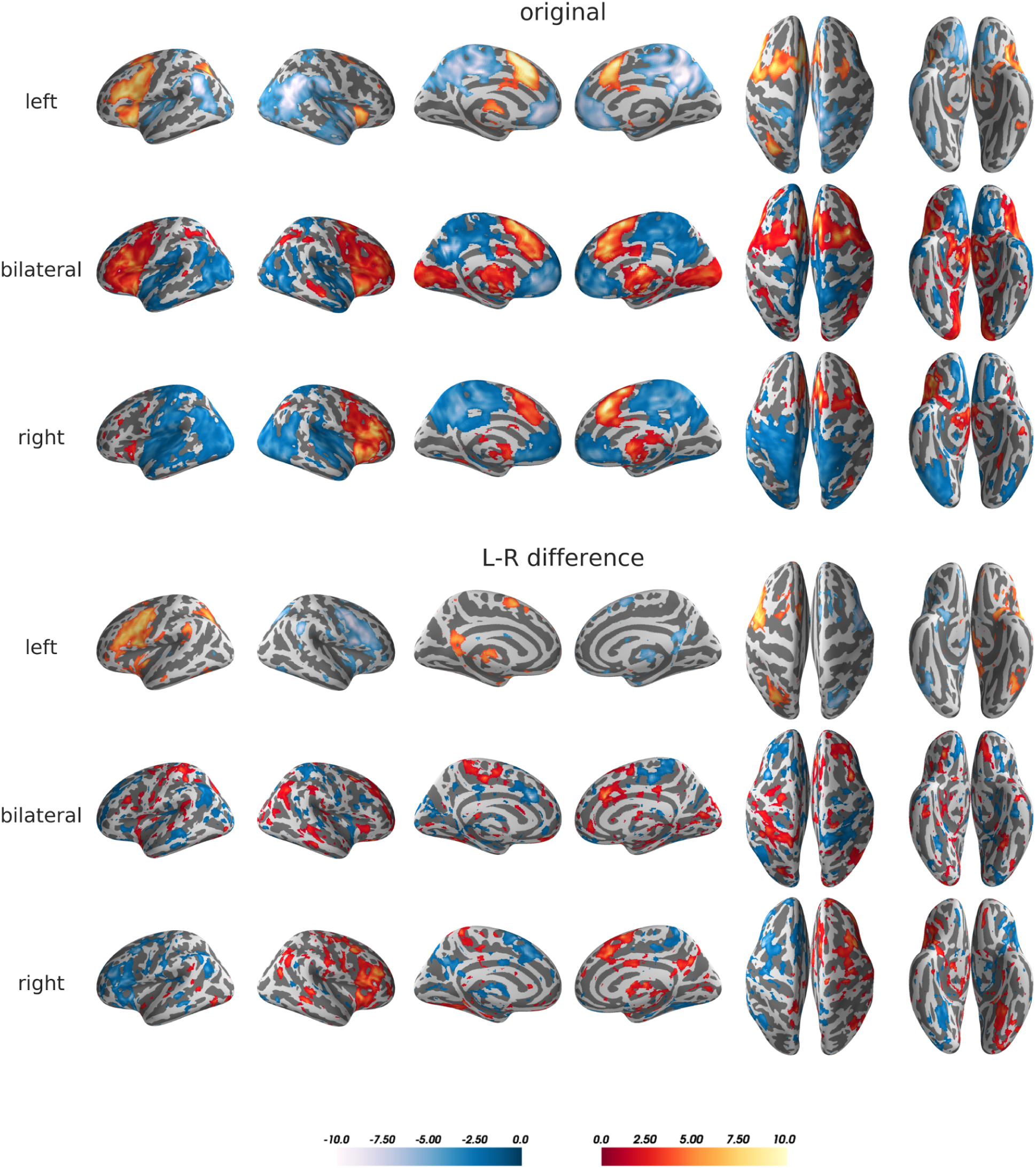
Maps of group results for the comparison between language task and rest. *original*: the task minus rest contrasts for each group were entered into mass-univariate one-sample t-tests on the group level. The alpha-level was adjusted for 1-beta=0.9 and d=0.8. Accordingly, thresholds of t=3.36 (left, N=39), t=1.23 (bilateral, N=10) and t=1.68, (right, N=14) were used, as computed using GPower 2.1. *L-R difference*: same as above, but each patient’s contrast image contained the left-right difference for each voxel, turning the maps of the left and right hemisphere into mirror-images of each other. Since the fsaverage template onto which the fMRI results were projected is not symmetrical, there appear to be slight asymmetries, although the maps are perfectly symmetrical in volume space. Full unthresholded maps of the group results are available on NeuroVault: identifiers.org/neurovault.collection:5132. The figure was created using pysurfer.

### 3.2 Volume-wise group analyses

The volume-template correlations for all patients in the training group are shown in Fig. 3A. Each of the three groups showed a characteristic pattern of correlations with the left-lateralized template. The group of typically left-lateralized patients showed positive correlations during task blocks and negative correlations during rest. The right-lateralized group showed the opposite pattern, with the two groups’ correlation patterns mirroring each other. The bilateral group’s correlation pattern was flatter and more closely distributed around zero, although it had the tendency to slightly mimic the left-lateralized group’s pattern of rise and fall. This general pattern was also present when averaging over one cycle of activity and rest (Fig. 3B).

**Figure 3.**
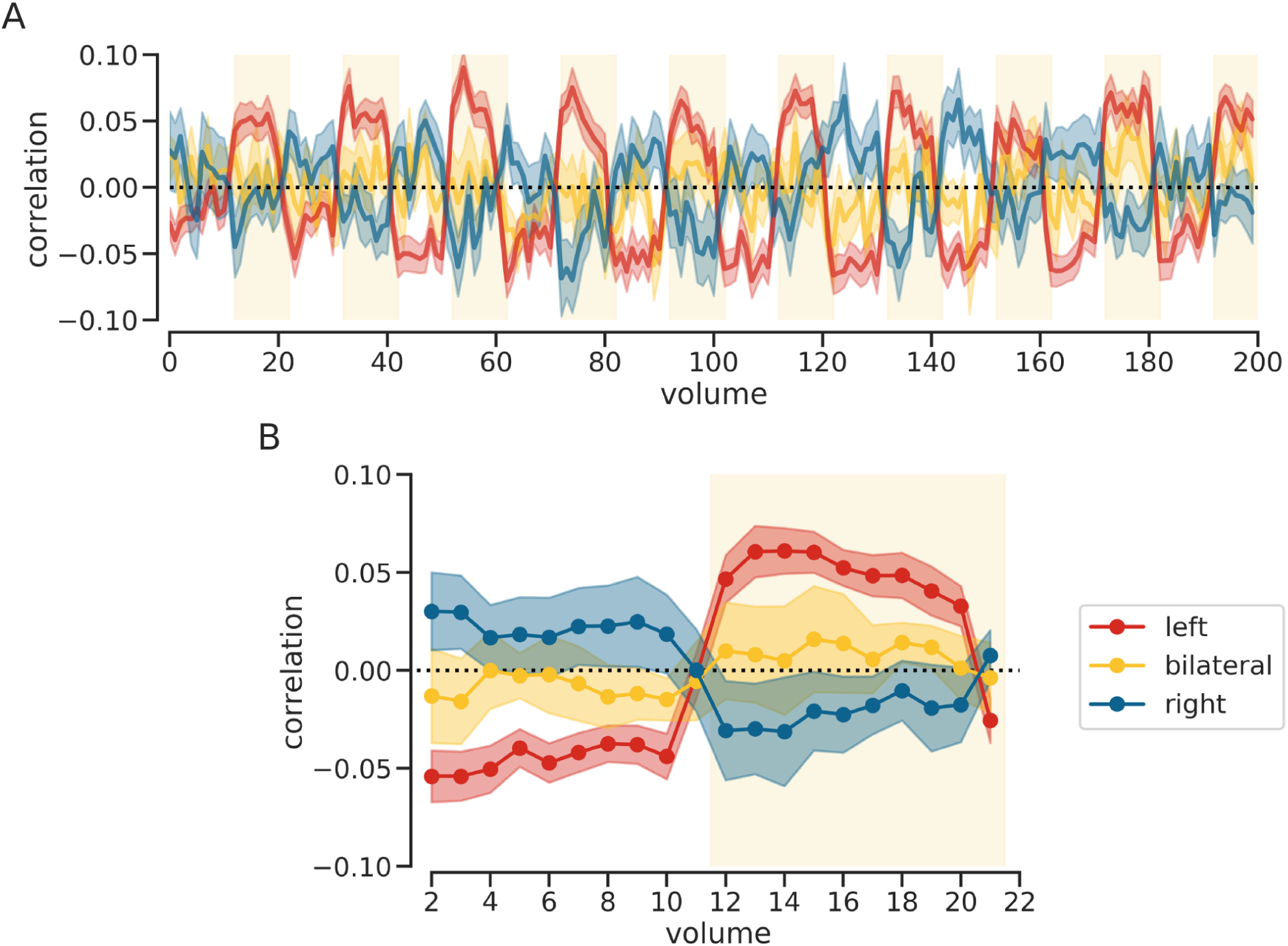
Volume-wise correlations on group level. Summary statistics were computed for each time point of the paradigm. Mean correlations (solid lines with strong color) with error bars (surrounding areas with light color) are shown for each lateralization group. A. Time course of correlations across the whole 200 volumes of the paradigm. The beige areas indicate periods when the language task was performed. Their onset is shifted by two volumes, in order to account for the expected lag in the hemodynamic response. Surrounding areas denote standard error. B. Time course of correlations averaged over one cycle of rest and activity. The first ten volumes (2 to 11) are rest, and the last ten volumes (12 to 21) are task. Surrounding areas denote 95% confidence intervals.

### 3.3 Training of classifier on single volumes

The distribution of correlations for all 12,600 volumes (200 x 63 patients) is shown in Fig. 4A. Correlations during rest were inverted for this step, so any positive correlations indicated left-lateralization while negative correlations indicated right-lateralization. As expected, volumes from left-and right-lateralized patients showed predominantly positive and negative values, respectively, while volumes of bilateral cases were centered around zero. These group differences were also reflected by non-overlapping means and 95% confidence intervals (Fig. 4B). The distribution of the classifier boundaries for all 63 folds of the leave-one-patient-out cross-validation scheme are shown as black vertical lines Fig. 4A and as individual points in Fig. 4C. These results indicate that there was a high degree of homogeneity across the different training folds. Overall, a volume was classified as bilateral when its correlation with the template was between -.015 and .036, classified as left-lateralized when the correlations were higher than .036 and classified as right-lateralized when they were smaller than -.015.

**Figure 4.**
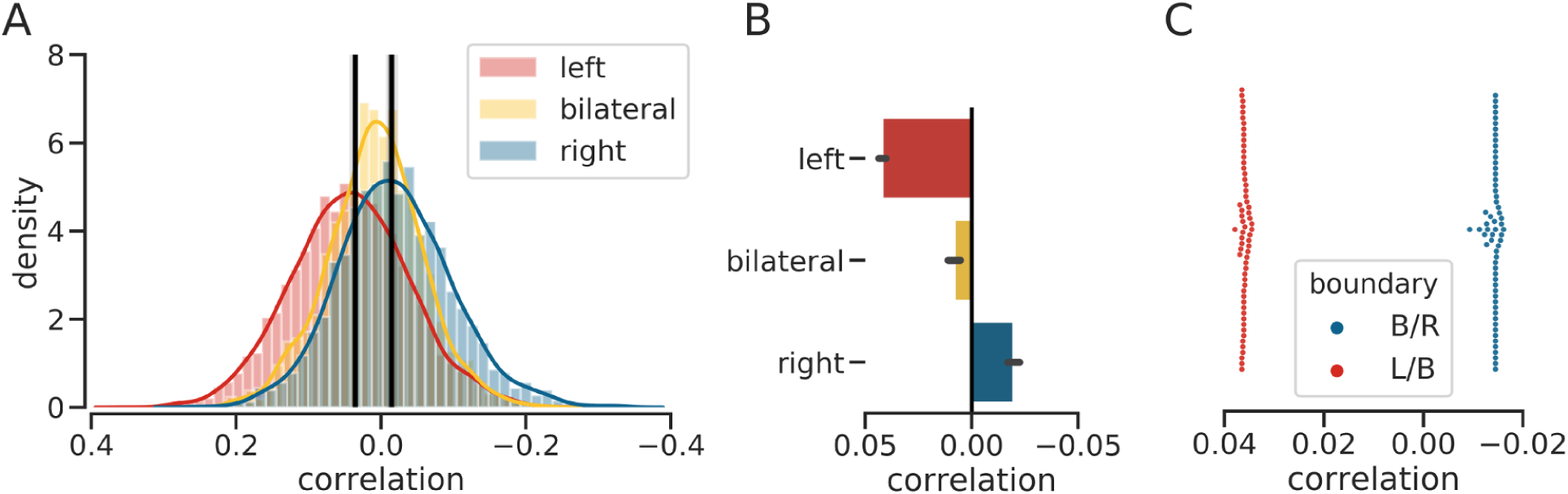
Training of classifier based on single volumes. A. For each of the three groups, a histogram with the distribution of correlations is shown. The class-boundaries for all folds of the leave-one-patient-out cross-validation are shown as black vertical lines. The lines are plotted with transparency, so that a stronger black indicates high overlap. B. For each of the three groups, mean and 95% confidence intervals of the correlations with the template are shown as summary statistics. Confidence intervals are tight as a total of 12,600 volumes were used as independent observations. C. Swarmplots illustrating the distribution of class-boundaries for all folds of the leave-one-patient-out cross-validation. These are the same data used to plot the black lines in part A. L/B= boundary between the left and bilateral class; B/R= boundary between the bilateral and right class. The x-axis has been inverted for all plots, so that correlations indicative of left-lateralization appear on the left side of the plots.

### 3.4 Evaluation of predictions

After the classifier boundaries from section 3.3 were applied to all of a patient’s 200 volumes, the patient had a triplet of values, which indicated the proportion of volumes which were classified as left-lateralized, bilateral or right-lateralized (Fig. 5). The patient’s type of language lateralization was then determined according to how most volumes had been classified. For two cases, a draw between two classes occurred and the predictions for these two cases were treated as incorrect.

**Figure 5.**
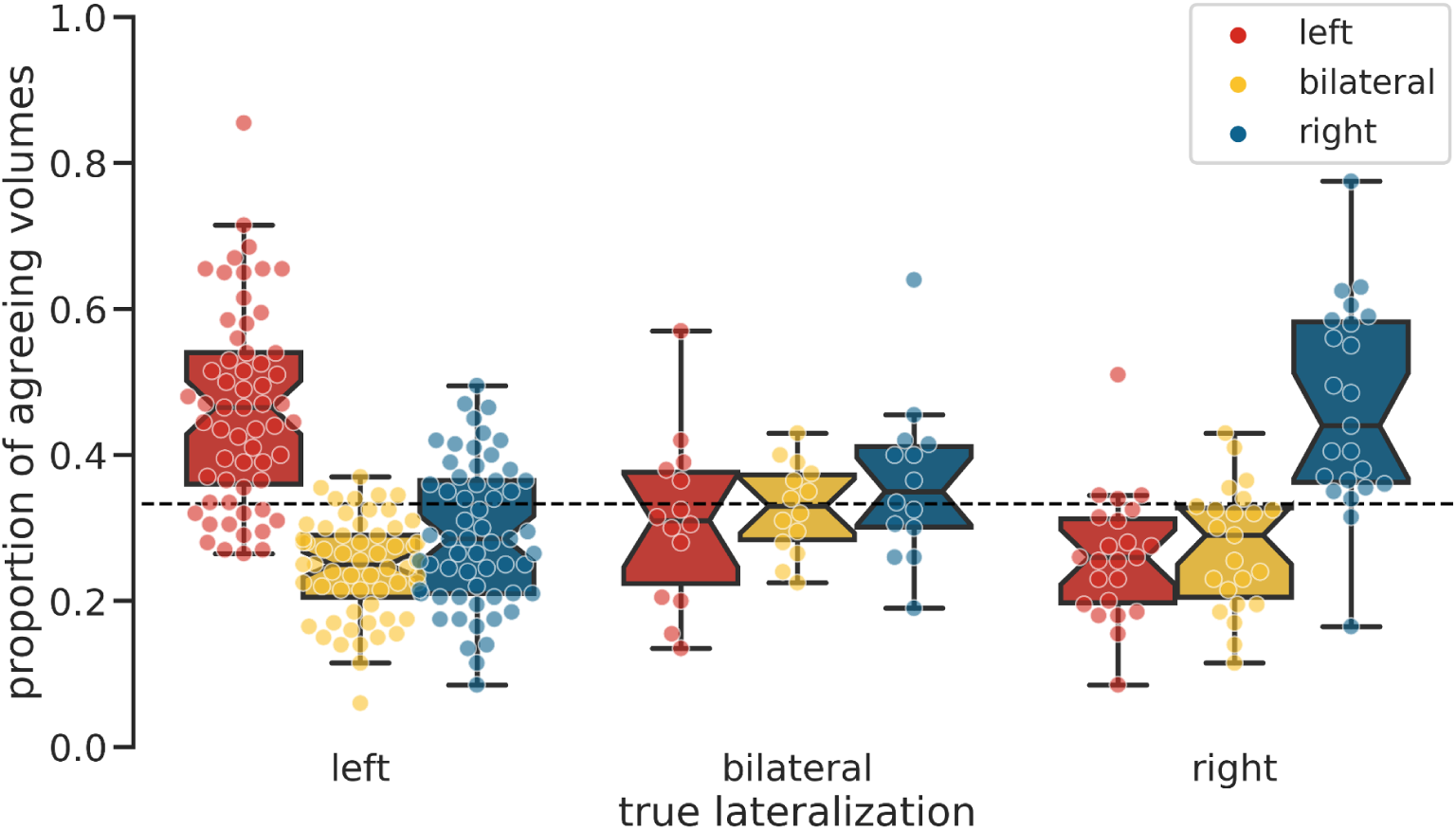
Proportion of volumes with the same predictions. For each patient, the proportion of volumes classified as left-lateralized, bilateral or right-lateralized are shown as dots. The color of the dots indicates which type of language lateralization the volumes were classified as. Notches of the boxplots indicate 95% confidence intervals. The dotted line at 1/3 indicates the expected proportion if all three classes were predicted equally often.

The classification accuracy for all preselected cases is shown in Fig. 6, at the point marked with (1). In total, 84% of cases were correctly classified as left-lateralized, bilateral or right-lateralized. The confusion matrix in Fig. 6B shows that while the predictions worked best for left-lateralized cases (97% correct), it was slightly lower for right-lateralized cases (79%) and much lower for bilateral ones (40%). However, if we considered only a division into typical (left) and atypical (bilateral and right) cases, the overall accuracy increased to 92%, with 97% accuracy for typical and 83%for atypical cases.

**Figure 6.**
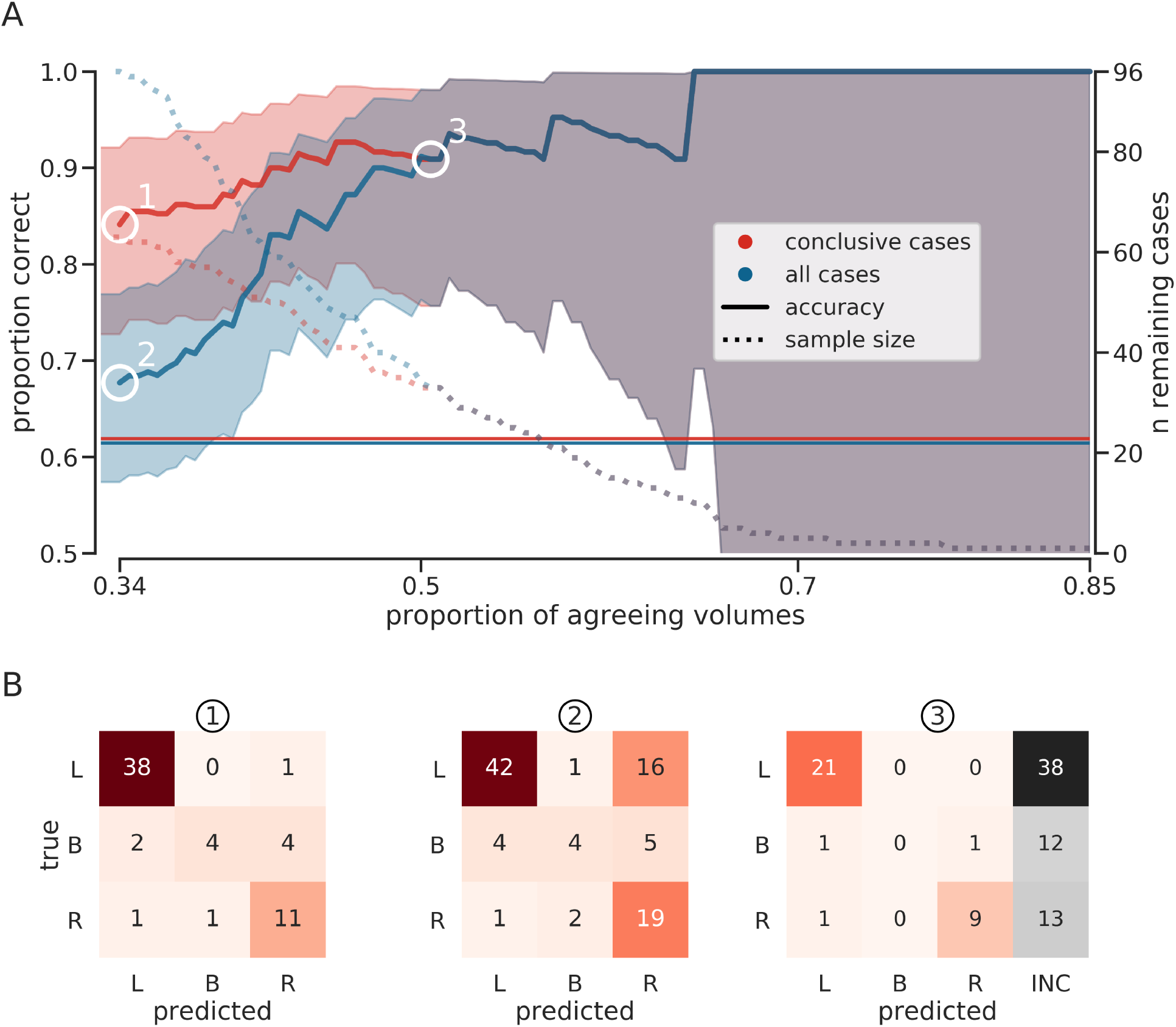
Evaluation of predictions for different cutoffs and group compositions. A. The accuracy of predictions and the number of remaining cases as a function of the cutoff used. The x-axis denotes the minimum proportion of agreeing volumes which has to be reached to make a decision. The y-axis on the left-hand side indicates the accuracy of the predictions of the Wada test results. Accuracies at different cutoffs (according to the proportion of agreeing volumes) are plotted as solid lines and their 95% confidence interval is plotted as a filled area. As the remaining sample size shrinks, the width of the confidence interval increases. The guessing rates for the two group compositions are indicated by solid horizontal lines (red line for the conclusive case group, blue line for the whole group). The y-axis on the right-hand side indicates the number of remaining cases, which are plotted as dashed lines. Three points are highlighted: (1) the accuracy at the lowest possible cutoff (34%) for cases preselected as conclusive; (2) the accuracy at the lowest possible cutoff for all cases, including ones which were deemed inconclusive; (3) the point of convergence beyond which it does not make a difference whether the data were preselected or not. B. Breakdown of the predictions at the three highlighted points using confusion matrices. Confusion matrices indicate the raw number of each type of prediction for each group. The two cases where a draw occurred were included as incorrect predictions when computing the accuracies in part A, but were omitted from the confusion matrices (1) and (2). L=left; B=bilateral; R=right; INC=inconclusive/no decision made.

The point marked (2) in Fig. 6 shows the accuracy when all 96 patients were included in the sample. In this case, overall accuracy dropped to 68% and the confidence interval included the 61% guessing rate (best guessing strategy being to always say left). However, Fig 6 also indicates that as the minimal number of volumes which needed to agree was increased, the accuracy for the group of patients who passed this cutoff also increased. When there were more than 43% of agreeing volumes, the accuracy for the sample with inconclusive patients reached over 84% correct, which was the initial accuracy for the preselected sample.

Furthermore, the point marked (3) shows that if there were at least 51% of agreeing volumes, both samples converged, meaning that all inconclusive patients were removed at this point, allowing to reach an accuracy of 91%. However, only 33 patients, roughly a third of the original sample, reached this criterion. Critically, no bilateral case remained after applying this cutoff, but many of the right-lateralized patients did; in fact, the ratio of right-lateralized to left-lateralized patients at this high cutoff was 10:21 (32% to 68%), comparable to the 22:59 (27% to 73%) in the initial full sample. At the tail end of Fig. 6, it can be seen that the predictions reached 100% accuracy at a cutoff of 65%, which a total of 10 patients were able to reach. Finally, the highest number of agreeing volumes for a single patient was 85.5%.

The proportion of volumes classified as left-lateralized, bilateral or right-lateralized for each patient is also shown in Fig. 7. As can be seen, the left- and right-lateralized patients reached high agreement rates of correctly classified volumes, while bilateral cases were classified correctly only by a tight margin. Fig. 7B illustrates the relationship of the proportion of agreeing volumes and the probability of making a correct decision using a logistic function. The increase in the logistic function showed a clear relationship of higher accuracy with an increasing number of agreeing volumes. The logistic function was used to derive an estimate of accuracy (y), given the number of agreeing volumes (x). This could allow for making a probabilistic statement about the language lateralization of an individual patient.

**Figure 7.**
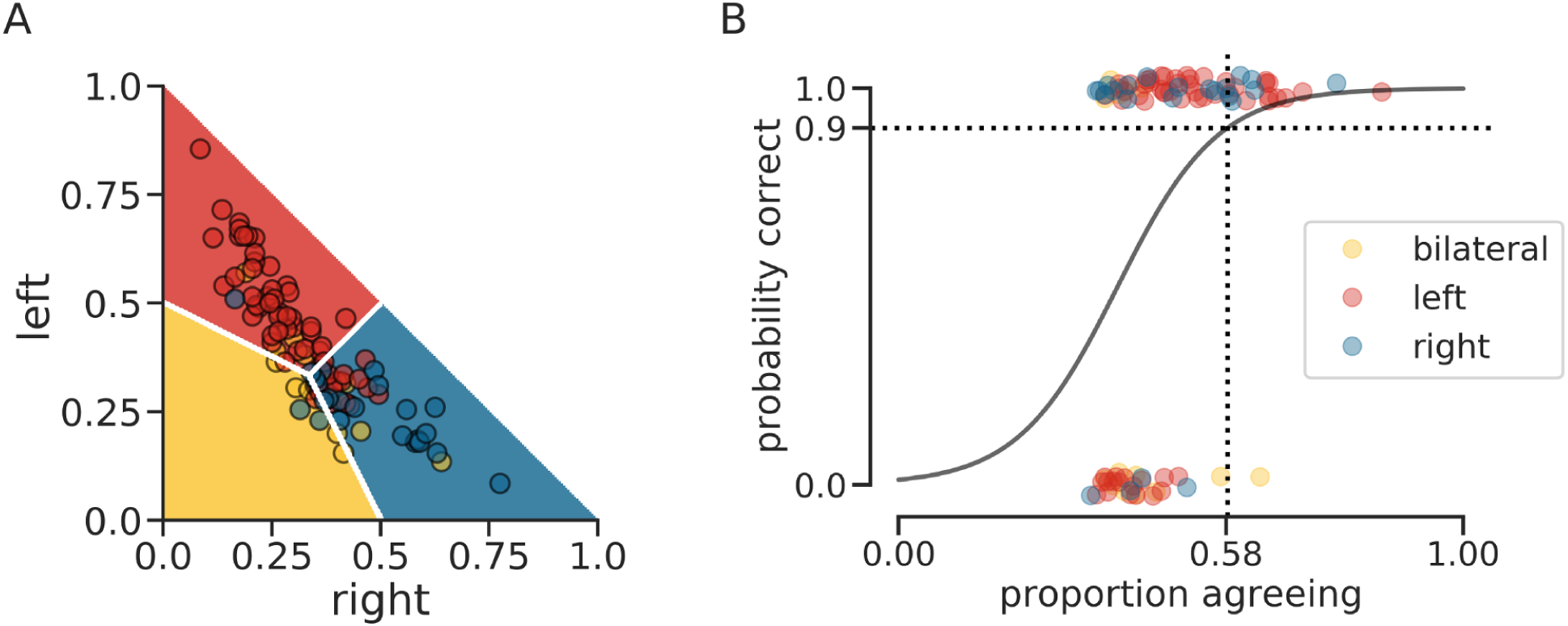
Distribution of agreeing volumes for all cases. Counts of agreeing volumes are shown for all N=96 cases. A. Proportion of volumes classified as left-lateralized, bilateral or right-lateralized for each patient. The color of each dot indicates the true lateralization, as determined by Wada testing. The position of each dot in the space indicates the proportion of volumes classified as belonging to each of the tree classes. Because the proportion of volumes always has to sum up to one, we can plot the three values of each patient in only two dimensions. Hence, the proportion of bilateral volumes corresponds to one minus the proportion of volumes classified as left and right (the remainder). The color of the background indicates which class most volumes fell into. B. Logistic regression fitted to the distribution of agreeing volumes for correctly and incorrectly classified patients. The numbers of agreeing volumes of patients classified correctly are shown at the top of the figure, those of patients incorrectly classified are shown at the bottom of the figure. The color of each dot indicates the true lateralization, as determined by Wada testing. The logistic regression tries to quantify the probability of being correct, depending on the number of agreeing volumes. As an example, the crossing dotted lines indicate how many agreeing volumes would be needed, according to the logistic regression, to reach an accuracy of 90%.

### 3.5 Application at the N=1 level

Fig. 8 illustrates how the analyses and classification schemes described above could be used for the prospective analysis of new cases. Except for the brain map in Fig. 8A, which illustrates the result of a classical voxel-wise t-test (task>rest); the remainder of Fig. 8 was created with a single command using the code on github.com/mwegrzyn/volume-wise-language. There, we also provide instructions and data which allow to generate such a figure, including all intermediate steps, for one healthy control. The figure illustrates how the classification of individual volumes could be visualized and how summary measures with attached probabilities could be provided for a single case. The time courses (Fig. 8B,C) allow for exploring the consistency of the patient’s data, while the summary statistics (Fig. 8D,E) permit making a simple winner-take-all decision. Finally, the value of the logistic function on the y-axis, given the patient’s number of agreeing volumes on the x-axis, could provide an estimate of the uncertainty associated with the decision.

**Figure 8.**
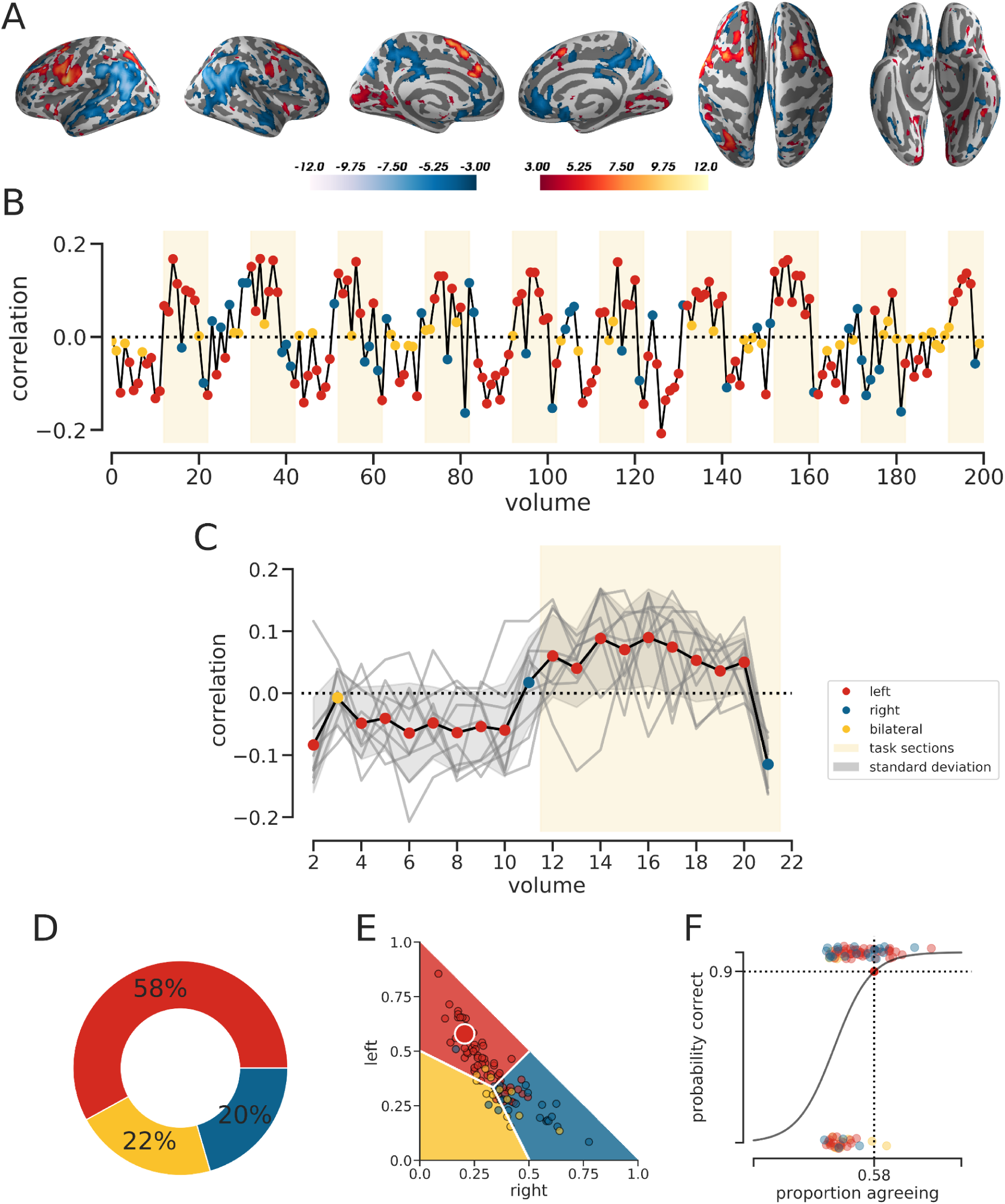
Example analysis for one patient. Illustration of how the analyses, fitted models and visualizations could be used on the level of N=1. A. Results of a t-test for the comparison of task > rest based on the patient’s original data. B. Volume-wise correlations across the whole paradigm, based on z-scored left-right images. Classifications of each volume are indicated by the color of the dots. Because the correlations have to be inverted during rest before they are classified, the same correlation value can be assigned different colors, depending on whether it occurs during task or during rest. C. Correlations averaged over one cycle of rest and activity. The gray lines are the time courses of each of the 10 individual cycles that constitute the mean. The gray background fill denotes the range of one standard deviation based on the patient’s data. D. Donut plot indicating the percentage of volumes classified as left-lateralized, bilateral or right-lateralized. E. Like Fig. 7A, but with the study sample plotted as small dots and the currently analyzed patient as a large dot. The color of the patient’s dot tells us which class was predicted by most of the patient’s volumes. F. Like Fig. 7B, but with the study sample plotted as small dots. The number of agreeing volumes of the patient is used to estimate the value of the logistic function at this point. Thus, given the number of agreeing volumes, we can read out the probability that the decision displayed in part E is correct.

## Discussion

In the present study, we explored how well single volumes of fMRI data allow for predicting language lateralization in patients with epilepsy. We used a correlation method to compare each volume of a patient’s fMRI time series with a template. Depending on the range into which the correlation fell, the volume was classified as indicative of left-, bilateral or right-lateralized language. To test the accuracy of the predictions, we used results from each patient’s Wada test as a gold standard. In a sample with around 60%typically lateralized cases, our method reached accuracies over 80% when trying to differentiate between left-, bilateral and right-lateralized cases. This corresponded to an accuracy of 92% when only considering the superordinate division into typical and atypical cases. Such a degree of accuracy is on par with previous work, for example the 91%accuracy reached in a sample with 70% typical cases in Woermann et al. (2003). Similarly, the meta-analysis by Dym et al. (2011) found accuracies of 88% for typical and 84% for atypical cases, while our analyses allowed us to reach accuracies of 97% correct for typical and 83% for atypical cases.

Extending these results, we showed that even if a large number of inconclusive cases is added to the dataset, those cases can be reliably filtered out by only using our main outcome metric. If at least 51% of volumes indicated the same lateralization, accuracies above 90% even for the three-class problem (left, bilateral, right) were reached, regardless of the initial composition of the dataset. Finally, all of the predictions were accurate for cases who had at least 65% agreeing volumes. This property of steadily increasing accuracy is important, as it has been acknowledged that successful language lateralization critically depends on data pre-selection (Wilke and Lidzba, 2007; Benjamin et al., 2017; Wegrzyn et al., 2019). In the current approach, as the number of agreeing volumes increased, so did the certainty with which a prediction could be made. Furthermore, this was true for both left- and right-lateralized patients, as the ratio of typical to atypical patients remained stable with more stringent cut-offs. This is an important extension of previous work, as language fMRI has been reported to be less reliable for patients with an atypical representation of language. The meta-analysis by Bauer et al. (2014) reported an accuracy of 94% for typical cases but a remarkably low accuracy of 51% for atypical ones. Based on these results, the authors recommended routine follow-up for patients with atypical language fMRI with a Wada test. Other authors have voiced similar concerns and recommended routinely repeating language fMRIs that indicate right dominance (Benjamin et al., 2018). Therefore, the present work is useful in showing that fMRI can identify right-lateralized cases with high confidence.

However, the same cannot be said of the bilateral cases, as our approach showed low sensitivity for this group: most bilateral cases were misclassified as left- or right-lateralized. Given that bilateral cases are notoriously difficult to classify (Rutten et al., 2002; Benke et al., 2006; Arora et al., 2009; Janecek et al., 2013), this low accuracy is not particularly out of line. More problematic is that the presented approach showed a bias against bilaterality. As can be seen in Fig. 7A, the model allowed only a few cases to have most of their volumes classified as bilateral. The few cases who were correctly classified as bilateral reached that class membership only by a tight margin. This is likely due to the model allowing only a narrow range of correlations to be classified as bilateral (Fig. 4). Although we used a balanced classification scheme, which should represent all groups equally, follow-up work needs to explore how to improve the representation of bilaterality. On a more speculative note, these findings might also be interpreted as indicating that bilaterality is not a steady state but rather a fluctuation of lateralization, constantly shifting between left- and right-hemisphere recruitment. Although it is beyond the scope of the present study to adequately address this question, our approach of analyzing the data in a volume-wise manner might provide a useful tool for addressing it in the future.

Analyzing the time courses volume-by-volume (Fig. 3) could also serve as a useful complement to inspecting only voxel-wise brain maps, as it allows for addressing new questions relevant to the clinical application of language fMRI. Being able to perform a group analysis for each of the 200 time points separately provided a unique perspective on the progression of the paradigm over time. The results show that there was no decrease in lateralizing information as the fMRI session progressed. Furthermore, there were no indicators that data quality deteriorated over time (e.g. due to increasing restlessness or fatigue). Hence, a ten minute session does not seem too long for the average patient. On the level of a single rest-activity cycle, it seemed that patients were performing the task over the entire 30-second length of a block. However, there seemed to be some decrease in lateralization and the drop in the HRF at the end (Fig. 3B) was slightly faster than the two volume shift expected by our model. Prospectively, such an analysis on the individual level could allow to identify patients who cannot come up with enough words to occupy themselves for the entire 30-seconds and might allow for selecting specific time ranges for re-analysis. For a paradigm that is performed covertly and without a measurable behavioral output, such an analysis of time course data could be particularly valuable. It could increase the diagnostic yield from otherwise inconclusive data and adjust for the differences in performance levels between patients.

Meanwhile, the current implementation of our approach is arguably most useful in the most clear-cut cases. While developing a method that can salvage more information from inconclusive data would be more desirable, the value of knowing which case can be diagnosed with particularly high certainty is valuable as well. Notably, the common implementations of the laterality index (Bradshaw et al., 2017a) do not have this property. Instead, they allow low-quality data to scatter across the whole spectrum of possible values, in often unpredictable ways (Wilke and Lidzba, 2007; Wegrzyn et al., 2019). In contrast, the present approach becomes increasingly more accurate as the number of agreeing volumes increases. By capitalizing on this property, it is possible to estimate the accuracy of each patient’s classification, instead of being able to make accuracy statements on the group level only.

Depending on one’s perspective, this approach can be seen as either a strength or a weakness. On the one hand, we want our diagnostic instrument to deliver clear categorical decisions, not fuzzy probabilistic statements. On the other hand, our decisions will be better if they explicitly consider uncertainty and embrace it as useful information (Gelman, 2018). The latter approach will enable the clinician to more efficiently integrate the fMRI results into a larger context. Such a context will include the results of other instruments, clinical variables such as handedness and age of onset (Berl et al., 2014), the possibility of ordering additional tests (Binder et al., 2008), the probability that repeating the measurement will give better data, the location of the planned resection (Labudda et al., 2018) and the consequences of making a wrong decision.

While the optimal integration of the current results into a larger framework is an important topic, the immediate question is how to further improve the accuracy of the present method. It is encouraging that, in the present sample, up to 171 out of 200 volumes of a patient’s fMRI session were correctly classified. This indicates that the employed correlation method can use each volume’s spatial information as a reliable fingerprint (Chang et al., 2015; Woo et al., 2017). Although correlations are robust and versatile (Hilborn and Mangel, 1997; Haxby, 2001), it would be interesting to know how our approach would compare to a volume-wise computation of laterality indices. Regarding the similarity of our approach with established methods, it is important to note that a single map of t-values can also capture the variability of the underlying data. Instead of counting volumes which resemble a certain pattern, counting the number of above-threshold voxels in a region of interest can also allow to make probabilistic predictions (Wegrzyn et al., 2019). Furthermore, the simplicity of our assumptions, while important to avoid overfitting and ensure transparency, is arguably detrimental in some regards. The simple boxcar function shifted by two volumes captured the signal reasonably well, but accounting for the gradual rise and fall of the hemodynamic response would certainly improve the handling of volumes during the transitory phases between activity and rest. Another improvement could be made at the later step of making a decision based on the number of a patient’s agreeing volumes. While using only the majority vote is straightforward, we could also use the specific combination of counts. That is, a case whose volumes were classified as being 51% bilateral, 48% left and 1% right should be treated differently than a case with volumes classified as 51% bilateral, 1% left and 48% right. Also, it is important to note that the presented approach, while using volume-wise information, cannot work when a patient provides only a single volume of data. The z-scoring of the voxels requires a full time series (the longer, the better) and a control condition against which the relative increase in language activity can be compared (Wegrzyn et al., 2018). As the z-scored values of each volume depend on the values of all the other volumes, it is necessary that the patient alternates between the language and comparison condition in accordance with the design. This also means that the approach cannot be applied to resting-state data or any design where a conventional contrast could not be computed as well. Finally, a limitation regarding the subsequent classification of volumes is that the method is only accurate if we compute the majority vote based on all volumes of a patient. The vote of a single volume will not be accurate once it is separated from the votes of the remaining volumes.

## Conclusion

In the present study, we explored how shifting from a voxel-wise to a volume-wise analysis of fMRI data can facilitate its clinical application on the level of N=1. We showed that aggregating all information within a single volume of fMRI data allowed us to predict Wada test results with high accuracy. Our outcome measure could be used to make categorical decisions, but could also serve as a measure of uncertainty. This property should allow to make more informed decisions on a single case basis. Hopefully, this way of translating fMRI data into clinically usable information will further increase the value it can offer to patients.

## Acknowledgments

Kirsten Labudda holds a Junior-Professorship at Bielefeld University endowed by the von Bodelschwinghsche Stiftungen Bethel. The funding source had no influence on the study’s design, data collection, analyses, interpretation, manuscript preparation and submission.

## Data Availability

fMRI group result maps are available on NeuroVault

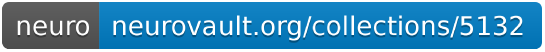

Code to reproduce the results and figures, as well as to recreate this manuscript, can be found on GitHub

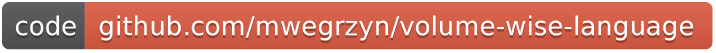

A sample dataset of a healthy control can be found on OpenNeuro

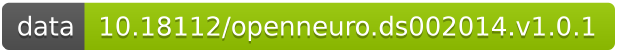

## Copyright

(C)2019 Wegrzyn et al. This article is distributed under the terms of the Creative Commons Attribution License, which permits unrestricted use, distribution, and reproduction in any medium, provided the original authors are credited.

